# Beyond Environmental Matching: Maternal Effects Depend on Offspring Environmental Complexity and Fitness Traits

**DOI:** 10.64898/2026.10.09.757982

**Authors:** Charlotte Carrier-Belleau, James A. Orr, Fanny Vermandele, Amy Bryan, Matthew Collins, Megan Glenn, Aisha Kyryk, Benjamin Spickett, Pepijn Luijckx, Jeremy J. Piggott

## Abstract

The conseuqences of maternal effects for offspring fitness can depend on the environmental conditions offspring encounter and the fitness traits considered. We used a transgenerational experiment in *Daphnia magna* to test how maternal exposure to warming and elevated salinity influenced offspring survival, reproduction and body size across matching, mismatching and multiple-stressor environments. Maternal exposure to salinity enhanced offspring survival under matching salinity, and benefits extended to mismatching environments. In contrast, maternal exposure to warming provided no survival benefit under matching warming and reduced survival in mismatching environments, while accelerating reproduction across contexts. Multiple-stressor conditions revealed pronounced maternal effects, while contrasting responses across survival, reproduction and body size showed that benefits to one fitness trait did not extend to others. Our findings demonstrate that environmental matching alone is insufficient to predict maternal effects, whose consequences depend on offspring environmental complexity and the fitness trait considered.

## 1. Introduction

Rapid environmental change increasingly exposes organisms to conditions that vary not only in magnitude, but also in their temporal predictability (Bernhardt *et al*. 2020; Bitter *et al*. 2021). Environmentally induced maternal effects, whereby environmental conditions experienced by mothers alter offspring phenotype and performance, represent a form of transgenerational plasticity (TGP) and may help organisms cope with environmental change by allowing offspring to respond before experiencing environmental conditions directly (Agrelius & Dudycha 2025). Hereafter, we use maternal effects to refer specifically to these environmentally induced responses. Critically, whether these effects are adaptive depends on how informative the maternal environment is about the conditions that offspring subsequently encounter (Kuijper & Hoyle 2015; Uller 2008). Theory predicts that adaptive transgenerational plasticity should be favoured when environments are sufficiently predictable across generations for parental experience to provide a reliable cue of future offspring conditions, whereas benefits should diminish when intergenerational correlations in environmental conditions are weak (Herman *et al*. 2014; Kuijper & Hoyle 2015; Leimar & McNamara 2015). Thus, adaptive transgenerational plasticity should depend not simply on maternal–offspring environmental matching, but on the intergenerational predictability underlying maternal cue reliability (Kuijper & Hoyle 2015; Uller *et al*. 2013).

Empirical tests of adaptive transgenerational plasticity have predominantly used matching designs, testing whether offspring perform better when they experience the same environmental conditions as their parents than when parental and offspring environments differ (Uller *et al*. 2013). Yet evidence for consistent fitness benefits of matching maternal and offspring environments remains weak (Burgess & Marshall 2011; Uller *et al*. 2013). More importantly, experimentally matching maternal and offspring environments does not establish that maternal conditions reliably predict offspring conditions in nature, a key requirement for anticipatory maternal effects to be adaptive (Kuijper & Hoyle 2015; Uller *et al*. 2013). Understanding when maternal effects are adaptive therefore requires distinguishing experimental environmental matching from the ecological context that determines the reliability of maternal cues.

Environmental predictability is not an intrinsic property of a stressor but emerges from its temporal dynamics within a particular ecological context. Environmental drivers such as salinity and temperature can vary over multiple temporal scales, meaning that the information maternal exposure provides about future offspring conditions will depend on local environmental dynamics. Consequently, comparisons among maternal environments can reveal whether their effects differ in magnitude and direction, but not whether one stressor constitutes an intrinsically more reliable cue than another. Cue reliability therefore provides an ecological framework for understanding why the benefits of maternal effects may vary across offspring environmental contexts.

Environmental context becomes especially important when offspring encounter multiple stressors, as interacting stressors increase the complexity of the conditions in which maternal effects are expressed. Transgenerational studies have commonly focused on maternal effects associated with a focal environmental driver, although maternal effects can also shape offspring responses to interacting stressors (Donelson *et al*. 2018; Sun *et al*. 2023; Tran *et al*. 2018; Vermandele *et al*. 2025). Because responses to multiple stressors cannot necessarily be predicted from responses to individual stressors, a maternally induced phenotype that enhances performance under one condition may provide little benefit or become costly when another stressor is present. Conversely, maternal exposure to one stressor may confer broader benefits if it increases tolerance across environmental contexts (Rodgers & Gomez Isaza 2023; Sun *et al*. 2023; Tran *et al*. 2018). We therefore expect the apparent benefit of a maternal effect to depend not only on whether maternal and offspring environments match, but also on the other stressors offspring encounter. Multiple-stressor environments may consequently expose benefits, costs and trade-offs of maternal effects that remain hidden in conventional single-stressor comparisons.

Whether a maternal effect appears adaptive should further depend on the fitness-related trait considered. Maternal effects on survival, growth and reproduction need not vary in parallel because resource allocation can generate life-history trade-offs (Marks & Lailvaux 2024; Marshall 2008). Benefits to one fitness trait may therefore be accompanied by costs to another, complicating their classification as uniformly adaptive or maladaptive (Hao *et al*. 2021; Marks & Lailvaux 2024). Evaluating multiple fitness-related traits is necessary to distinguish broadly beneficial maternal effects from shifts in life-history allocation that improve one component of performance at the expense of another (Marks & Lailvaux 2024).

Here, we used a transgenerational experiment in *Daphnia magna* to test whether the consequences of maternal effects depend on environmental matching, offspring environmental complexity and the fitness-related trait considered. The genotype used in our experiment originates from a coastal rock pool, where both salinity and temperature represent ecologically relevant sources of environmental variation. Mothers were exposed to elevated salinity and warming, individually and in combination, and offspring from control-, salinity– and warming-exposed mothers were subsequently exposed to control conditions, salinity, warming or their combination. We quantified survival, reproductive timing and output, and body size across matched, mismatched and multiple-stressor conditions. Specifically, we predicted that (1) maternal exposure to a given stressor would benefit offspring subsequently exposed to the same stressor if maternal effects are anticipatory; (2) these benefits would weaken or disappear when maternal and offspring environments mismatched, while multiple-stressor environments would reveal whether maternal effects remain stressor-specific or extend across environmental contexts; and (3) maternal effects would differ among survival, reproduction and body size, revealing potential life-history trade-offs. Together, this design tests whether environmental matching consistently predicts beneficial maternal effects, or whether their consequences instead depend on offspring environmental complexity and the fitness trait considered.

## 2. Material and Methods

### 2.1. Preparation of parental generation from stock culture and acclimation under control conditions

*Daphnia magna* is a cyclically parthenogenetic zooplankton with a short generation time and rapid reproduction, enabling experimental manipulation across generations while controlling for genetic variation through clonal reproduction. Thirty-six adult females from the clonal line Fi-Oer-3-3, (Finland: 59.7886° N, 23.1741° E) originally isolated from a rock pool and maintained for >50 generations under standardized laboratory conditions, were cultured for three weeks before the experiment (stock culture; Fig. 1A). To obtain sufficient individuals for the experiment, groups of 10–12 females were maintained in three 400-mL glass microcosms containing 200 mL artificial *Daphnia* medium (ADaM; Klüttgen *et al*. 1994; baseline salinity: 0.33 g L□^1^) at 20°C under a 16:8 h light:dark photoperiod and fed *ad libitum* with batch-cultured *Acutodesmus obliquus* (Oliveira *et al*. 2021). Algae were cultured at 20°C in WC medium (Kilham *et al*. 1998) under nutrient– and light-saturated conditions, with 40% medium replacement three times weekly; while *Daphnia* medium was renewed twice weekly. Three days before the acclimation phase, adults were transferred to fresh microcosms, and juveniles produced during the subsequent 72 h were collected and sexed under a dissecting microscope (OM2344, Omano, USA). Juvenile females (1–3 days old) constituted the parental generation (P) and were individually acclimated for 16 days under control conditions (0.33 g L□^1^ salt, 20°C) in 144 100-mL glass microcosms (Fig. 1B). Microcosms were distributed among six trays (24 microcosms per tray) across three temperature-controlled water baths (two trays per bath). Individuals were transferred to fresh medium twice weekly, with offspring removed. Animals were fed three times weekly at concentrations increasing from 5 × 10□ to 1.2 × 10□ algal cells mL□^1^ by day 7. At the end of acclimation, offspring produced during the final three days were collected and sexed, and females were pooled for <3 h before assignment to their respective experimental treatments. These females constituted the F1 generation used for maternal exposure.

**Figure 1.**
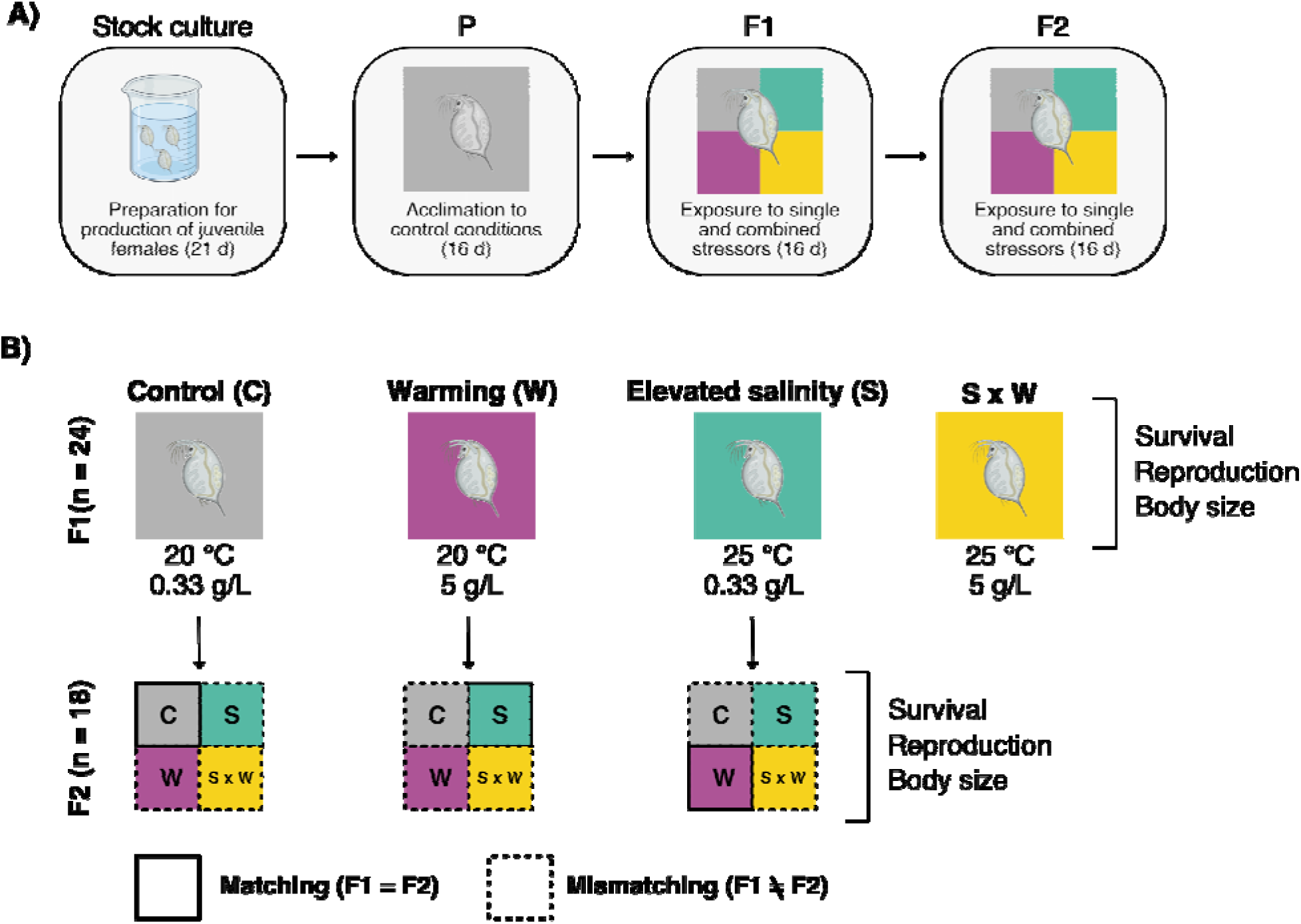
Experimental design illustrating transgenerational exposure to single and combined environmental stressors. (A) Experimental timeline showing acclimation under control conditions of the parental generation (P), followed by experimental exposure of mothers (F1) and offspring (F2) to warming, elevated salinity and combined stressors. (B) Factorial design depicting maternal (F1) exposure to elevated salinity (S; green), warming (W; pink), and combined stressors (S × W; yellow), followed by offspring (F2) exposure to all four environmental conditions. Solid lines indicate matching maternal–offspring environments, whereas dashed lines indicate mismatching environments between the two generations.

### 2.2. Maternal (F1) and offspring (F2) experimental exposures

To assess individual and combined effects of elevated salinity and warming across generations, parental (F1) females and their offspring (F2) were subsequently exposed for 16 days to four treatments, using a full factorial design: control (0.33 g L□^1^ salt, 20°C), elevated salinity (5 g L□^1^, 20°C), warming (0.33 g L□^1^, 25°C), or combined salinity and warming (5 g L□^1^, 25°C; Fig. 1B). Treatment levels were selected based on previous studies showing that salinities of approximately 4–6 g L□^1^ and temperatures of 25°C represent physiologically relevant stress conditions for *D. magna* while remaining within its tolerance range (Im *et al*. 2020; Martínez-Jerónimo & Martínez-Jerónimo 2007). Elevated salinity was achieved by supplementing ADaM with synthetic sea salt (Instant Ocean, Aquarium Systems).

For F1, each treatment comprised 24 replicate microcosms distributed across 12 trays and six water baths (three at each temperature; Fig. 1B). After 16 days, female offspring (F2) from control-, salinity– and warming-exposed mothers (F1) were collected within 12 h of birth and pooled by maternal treatment for <3 h before F2 exposure. Due to high mortality, offspring from F1 mothers exposed to the combined conditions were unavailable in sufficient numbers to start the F2 exposure.

F2 females from each maternal treatment (control, salinity or warming) were exposed to the same four treatments, yielding 12 maternal × offspring environments (Fig. 1B). Each combination comprised 18 replicate microcosms distributed across six trays and three water baths at the corresponding offspring temperature. Within trays, maternal origins (i.e. control, warming, elevated salinity) were arranged across columns and offspring treatments (i.e. control, warming, elevated salinity, warming x elevated salinity) across rows. During both generations, individuals were transferred to fresh treatment medium twice weekly and fed three times weekly with 1 mL algae, increasing from 5 × 10□ to 1.2 × 10□ cells mL□^1^ by day 7. Tray positions within water baths were rotated after each transfer to minimize positional effects. Temperature, salinity and conductivity were measured twice weekly before transfers in one jar per tray within each water bath using a portable multi-meter (HQ2200, Hach). The measured values are presented in Table S1.

### 2.3. Measurements of fitness and life-history traits

Survival and reproduction were monitored three times weekly throughout F1 and F2 exposures. Survival was expressed as survival probability, while reproduction was quantified as cumulative reproduction probability (the probability of having reproduced by a given day) and total offspring production. At the end of each 16-day exposure, body length (head to base of the tail spine) was measured in surviving individuals using a stereomicroscope with an ocular micrometer (Olympus SZ61). Maternal (F1) survival, reproduction and body size responses to the experimental treatments are presented in Figure S1.

### 2.4. Statistical analyses

Statistical analyses were conducted using Bayesian hierarchical models implemented in R (version 4.5.2) using the brms package (Bürkner 2017), which interfaces with the probabilistic programming language Stan (Carpenter *et al*. 2017). Bayesian models allowed us to incorporate weakly informative priors in a transparent way, which constrained parameter estimates to biologically plausible ranges and enhanced convergence of our relatively complex models. Working with posterior distributions also allowed us to communicate contrasts between treatments, effect sizes of treatments and uncertainty in an intuitive way (McElreath 2020). Following the factorial experimental design, temperature in F2, salinity in F2, environment of F1, and their interactions (two-way and three-way) were included as predictors of parameters in all models of the responses of F2 individuals. Additionally, random intercepts for experimental tray nested within bath were included to account for the hierarchical design and potential non-independence among individuals from the same experimental groups. Models were fit using four MCMC chains, with 2000 iterations per chain (including 1000 warm-up iterations). Sampling performance was assessed using effective sample sizes, R-hat statistics, and examination of posterior trace plots. Posterior predictive checks were used to evaluate model fit by comparing observed data with predictions generated from the posterior distribution. Prior predictive checks were also used to ensure that our priors generated biologically plausible predictions but did not strongly bias our posterior distributions.

Time-to-event models were used to estimate treatment effects on survival and time to reproduction. These data were right-censored because individuals that survived until the end of the experiment were only known to have survived for at least 16 days (Fox 2001; McElreath 2020). Similarly, individuals that survived but did not reproduce during the experimental period were only known to have a time to reproduction of at least 16 days. We therefore used survival models with censoring structures rather than treating these observations as exact event times. We used Weibull models for these two responses as they are flexible in the sense that their hazard (instantaneous rate of the event occurring, conditional on the event having not occurred so far) can change (monotonically) over time, but they remain easy to interpret with just two parameters (Fox 2001). The “scale” parameter determines the overall time-to-event, while the “shape” parameter determines how the hazard changes over time. Weakly informative priors were assigned to both Weibull parameters. For the intercepts of the scale parameter (on the log scale), we used a normal distribution with mean 3 and standard deviation 1 for the survival model and with mean 2 and standard deviation 0.5 for the reproduction model. These priors focus the prior probability mass around baseline survival times of ∼20 days (with a 95% prior interval of approximately 3 to 150 days) and baseline times to reproduction of ∼7.5 days (with a 95% prior interval of approximately 2 to 20 days). These priors therefore penalise extremely long survival and time-to-reproduction times while remaining sufficiently weak that posterior inference was primarily informed by the observed data. Regression coefficients for treatment effects on the scale parameter were assigned normal priors with mean 0 and standard deviation 1.5, allowing both strong positive and negative treatment effects. For the shape parameter (on the log scale), the intercept and treatment coefficients were assigned normal priors with mean 0 and standard deviation 1.5. This corresponds to a broad prior allowing increasing, constant, or decreasing hazards over time. Exponential priors with rate 1 were assigned to the standard deviations of random effects, providing regularisation while allowing variation among experimental groups and enabling partial pooling.

Total offspring production of reproducing daphnia and body size of surviving daphnia were modelled using standard Bayesian hierarchical generalised linear models (McElreath 2020). These analyses were conditional on reproduction and survival, respectively, and therefore quantify treatment effects on the reproductive output of reproducing individuals and the body size of surviving individuals, rather than population-level responses. Total offspring production was modelled using a Poisson error distribution with a log link, while body size was modelled using a Gaussian error distribution with an identity link. For both models, treatment coefficients were assigned weak normal priors with mean 0 and standard deviation 2. Intercepts were assigned weak normal priors with mean 3 and standard deviation 2 for the offspring model and mean 4 and standard deviation 2 for the body size model, therefore centred on biologically plausbile values but remaining extremely broad. Full posterior contrasts for all response variables are provided in Tables S2-S5.

## 3. Results

Maternal (F1) survival, cumulative reproduction probability, total offspring production and body size across experimental environments are shown descriptively in Fig. S1 to provide context for maternal performance. Maternal environments subsequently generated contrasting transgenerational effects across offspring environments and fitness-related traits.

### 3.1 Maternal exposure to warming reduces survival but accelerates offspring life histories

Maternal exposure to warming generated contrasting transgenerational effects across fitness traits. It reduced offspring survival under mismatching conditions but had no detectable effect on survival under matching warming, while accelerating reproduction across most F2 environments (Figures 2–4).

**Figure 2.**
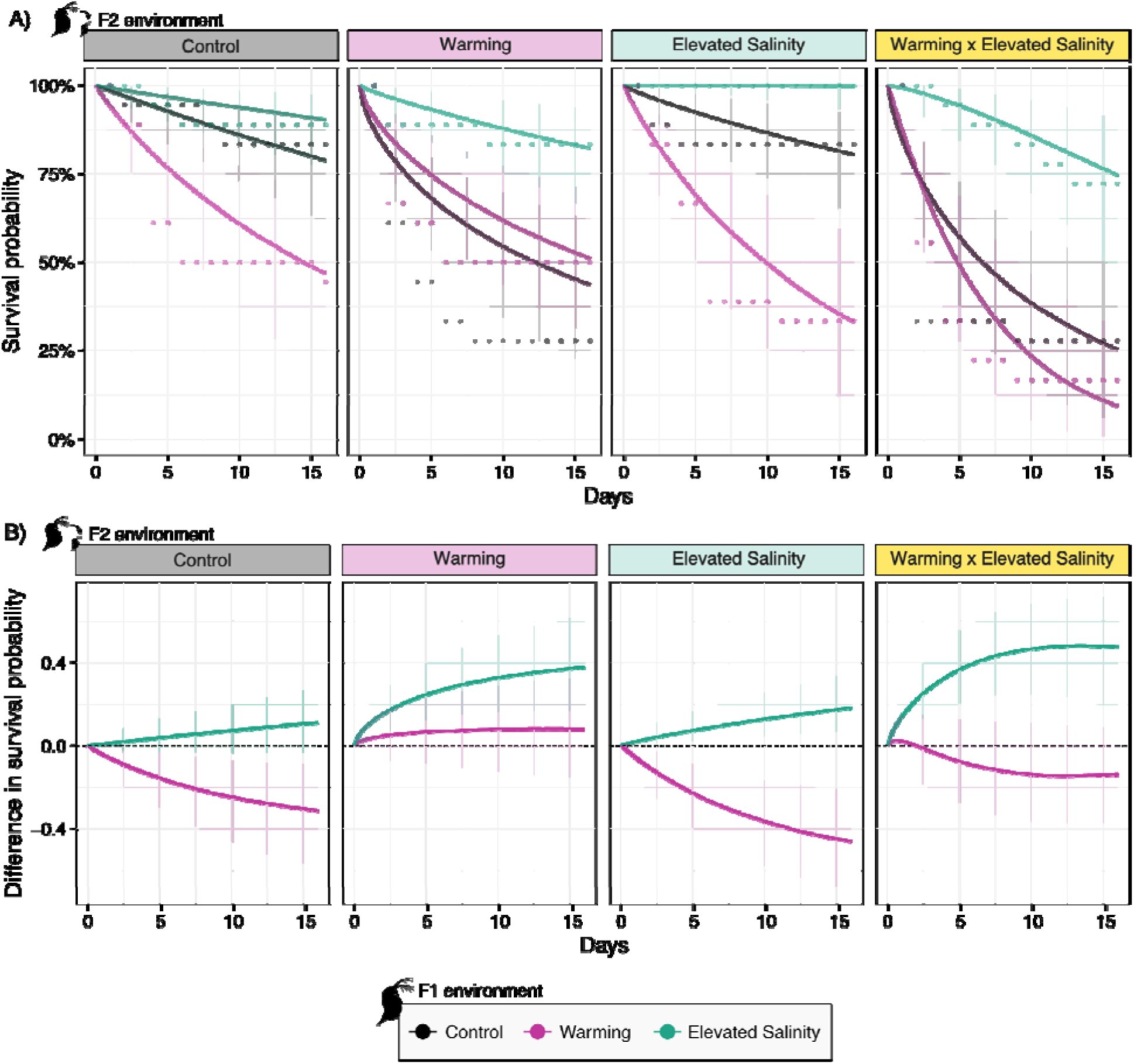
Transgenerational effects of maternal exposure to warming (pink), elevated salinity (green) and under control conditions (black) on offspring survival across single and combined stressor environments. Top panels (A) show model fits for offspring survival trajectories across treatments with empirical observations, while bottom panels (B) illustrate the magnitude and direction of maternal effects relative to controls. Lines represent median posterior predictions, and shaded areas represent 90% credible intervals.

**Figure 3.**
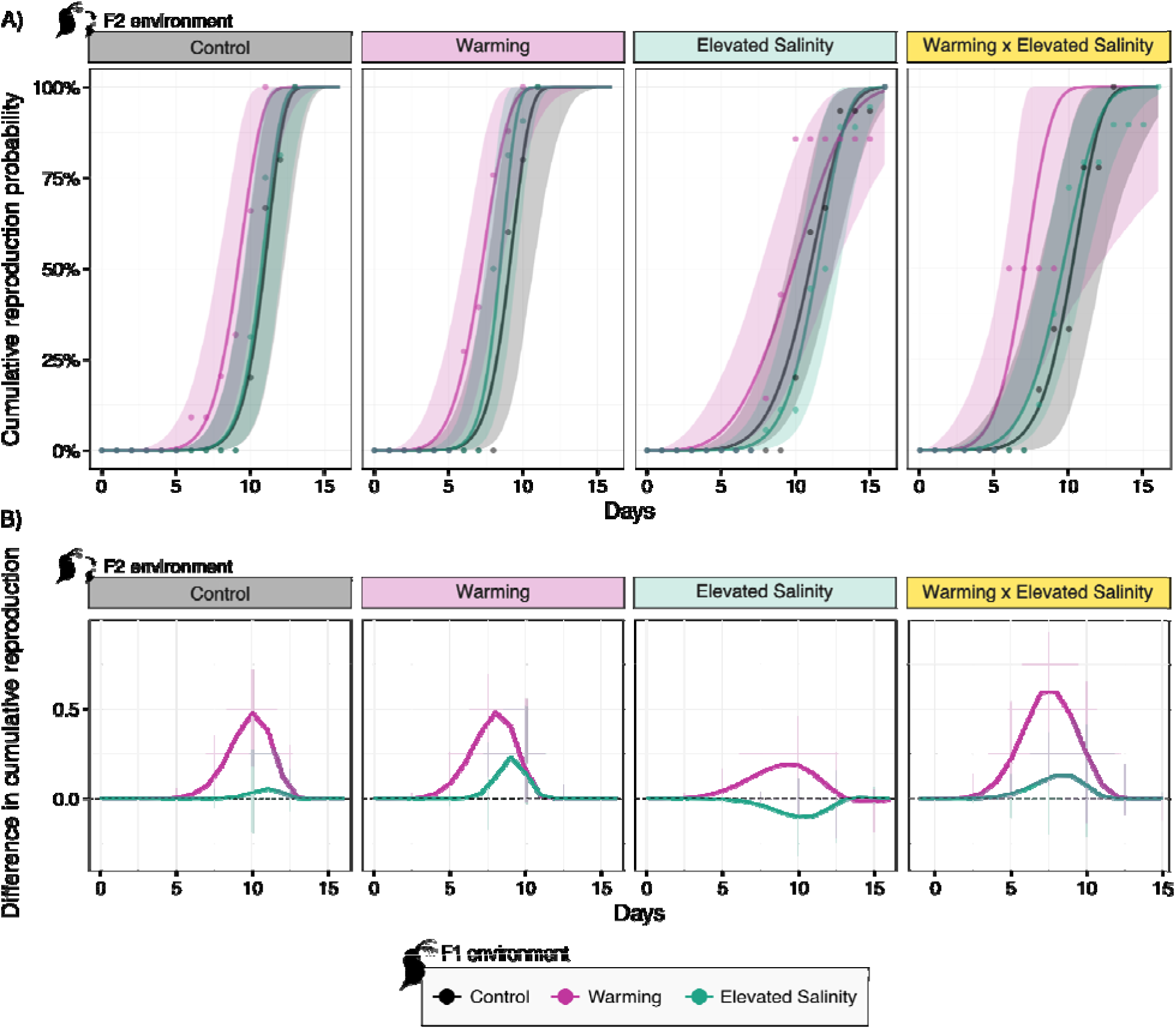
Transgenerational effects of maternal exposure to warming (pink), elevated salinity (green) and under control conditions (black) on offspring cumulative reproduction across single and combined stressor environments. Top panels (A) show model fits for offspring cumulative reproduction probability across treatments with empirical observations, while bottom panels (B) illustrate the magnitude and direction of maternal effects relative to controls. Lines represent median posterior predictions, and shaded areas represent 90% credible intervals.

**Figure 4.**
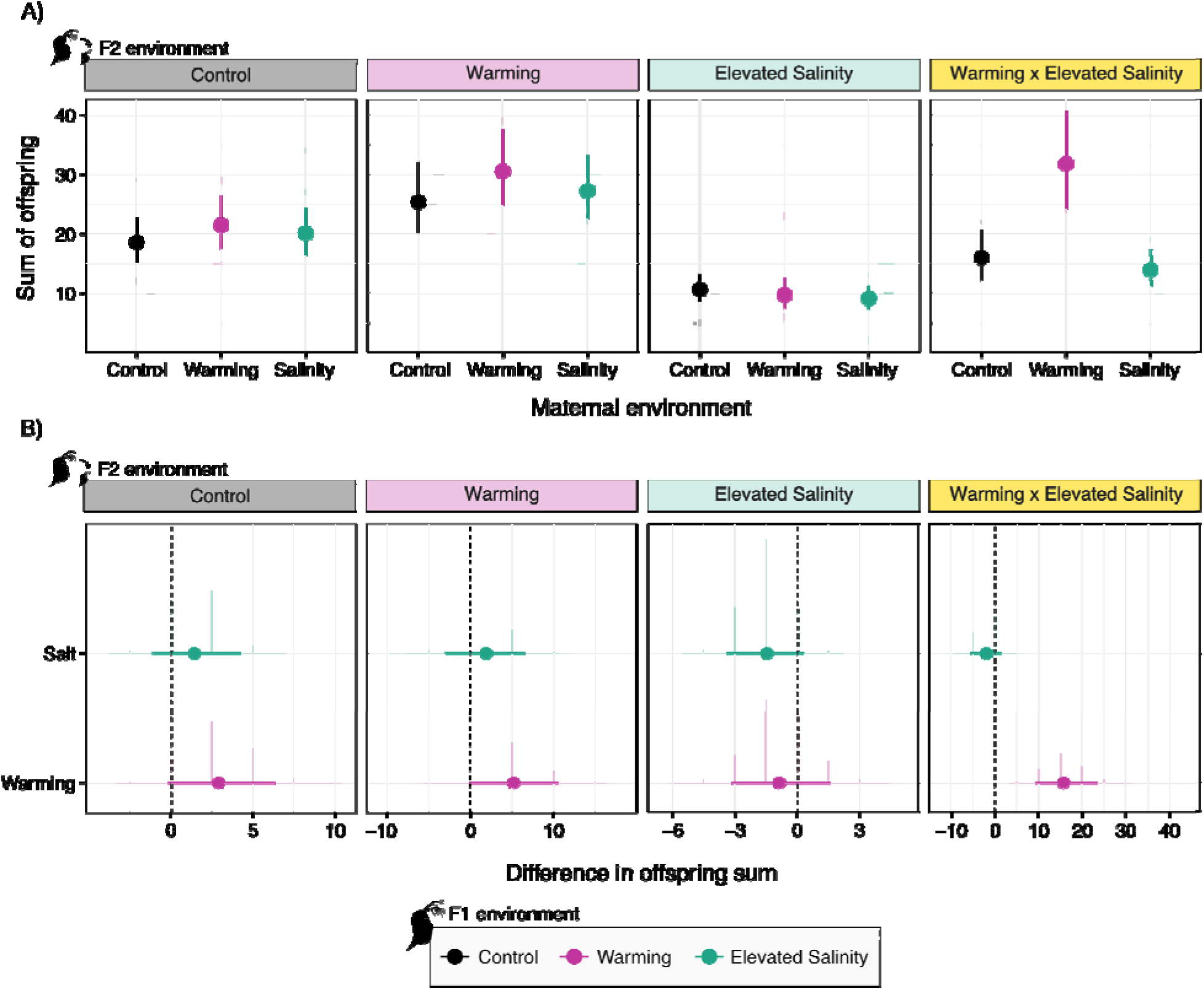
Transgenerational effects of maternal exposure to warming (pink), elevated salinity (green) and under control conditions (black) on offspring total reproduction across single and combined stressor environments. Top panels (A) show model fits for offspring total reproduction (i.e. sum of offspring) across treatments with empirical observations, while bottom panels (B) illustrate the magnitude and direction of maternal effects relative to controls. The full circle represent median posterior predictions, lines represent 90% credible intervals, and shaded areas represent full posteriors.

Maternal exposure to warming imposed time-increasing survival costs, with the strongest negative effects when offspring experienced environments differing from the maternal environment. Under F2 control conditions, survival deficits relative to control maternal offspring increased from –0.217 at day 8 (90% CI –0.407 to –0.057) to –0.313 at day 16 (90% CI –0.582 to –0.090; Figure 2B). The strongest survival costs occurred under F2 exposure to elevated salinity, where contrasts reached –0.319 at day 8 (90% CI –0.523 to –0.144) and –0.462 at day 16 (90% CI –0.692 to –0.214). Under combined warming and salinity, contrasts were consistently negative but 90% credible intervals overlapped zero throughout the experiment, providing no clear evidence of a maternal warming effect on survival. When offspring experienced matching warming conditions, survival differences were small and uncertain (e.g. day 8: +0.074, 90% CI –0.111 to +0.263), providing no evidence of a survival benefit from maternal warming (Figure 2B).

Maternal exposure to warming accelerated reproduction across most F2 environments (Figure 3A). At day 8, cumulative reproduction probability was higher in offspring under control (+0.185, 90% CI +0.027 to +0.443), warming (+0.482, +0.137 to +0.753), and combined warming–salinity conditions (+0.592, +0.085 to +0.936; Figure 3B). This acceleration translated into higher total offspring production under combined stressors (day 16: +15.683, 90% CI +9.326 to +23.313; Figure 4), while the increase under warming alone was more uncertain (+5.164, –0.100 to +10.472; 94.6% posterior probability of a positive effect). Maternal exposure to warming did not impact cumulative reproduction or total fecundity when offspring experienced elevated salinity. Maternal exposure to warming had no detectable effect on offspring body size in any F2 environment.

### 3.2 Maternal exposure to elevated salinity enhances survival with cross-trait costs

Maternal exposure to elevated salinity generated strong survival benefits across stressful F2 environments (Figure 2), while effects on reproduction (Figures 3,4) and body size (Figure 5) were weaker. Under matching F1-F2 elevated salinity conditions, offspring showed clear and time-increasing survival advantages (e.g. day 8, median difference +0.107, 90% CI +0.034 to +0.221; day 16, +0.182, 90% CI +0.065 to +0.348; Figure 2B). Survival reached 100% for offspring in the matching elevated salinity maternal-offspring environment, potentially limiting the scope for additional maternal buffering to be expressed. Importantly, survival benefits extended beyond the matching environment. For offspring exposed to warming, contrasts increased from +0.302 at day 8 (90% CI +0.150 to +0.486) to +0.378 at day 16 (+0.167 to +0.621). The largest observed differences occurred for offspring under combined warming and salinity, reaching +0.440 at day 8 (90% CI +0.242 to +0.653) and +0.475 at day 16 (+0.175 to +0.716). In contrast, maternal salinity had no clear survival effect on offspring reared under control conditions (day 8, +0.059, 90% CI –0.036 to +0.172).

**Figure 5.**
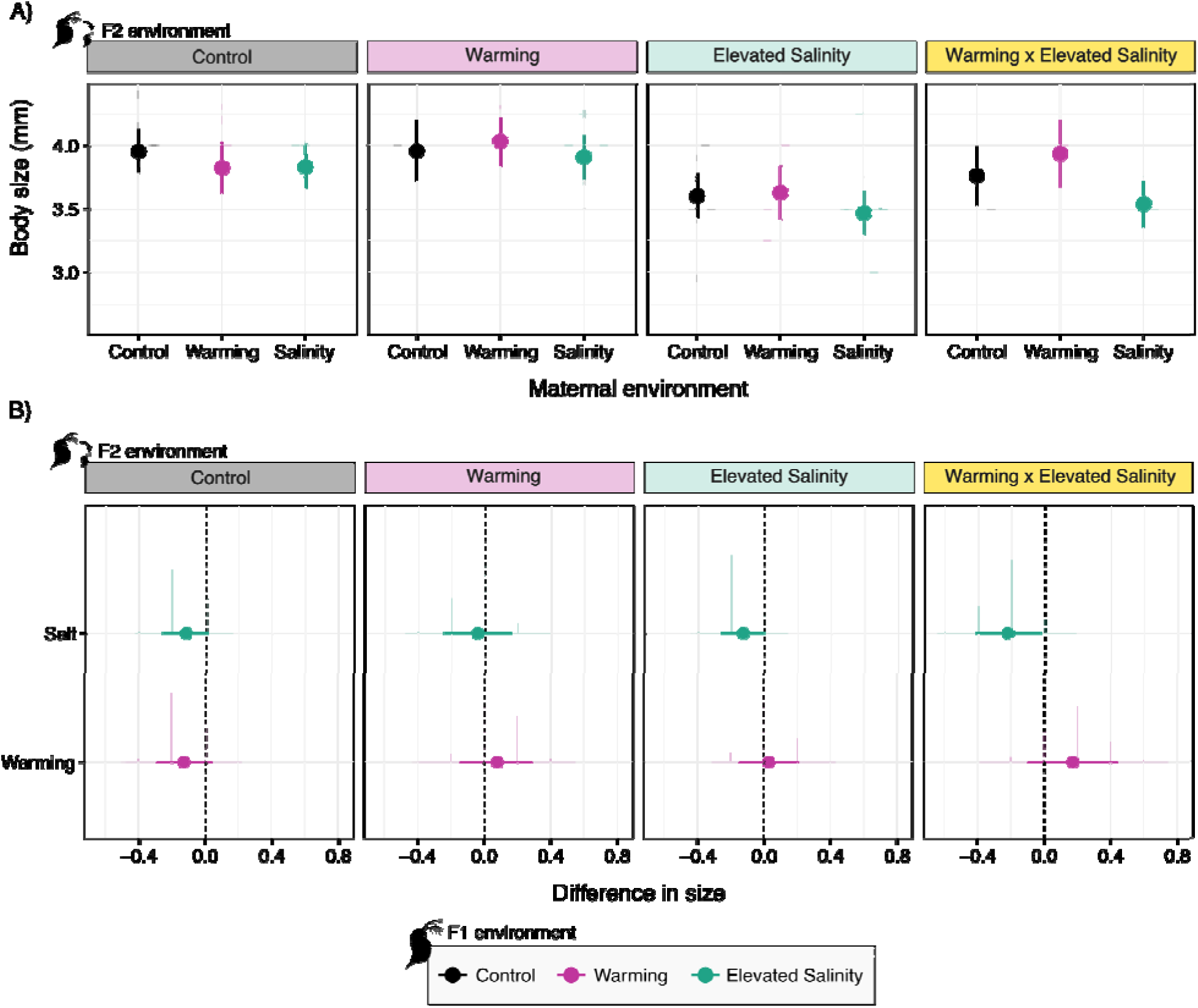
Transgenerational effects of maternal exposure to warming (pink), elevated salinity (green) and under control conditions (black) on offspring body size across single and combined stressor environments. Top panels (A) show model fits for offspring body size across treatment with empirical observations, while bottom panels (B) illustrate the magnitude and direction of maternal effects relative to controls. The full circles represent median posterior predictions, line represent 90% credible intervals, and shaded areas represent full posteriors.

Maternal exposure to elevated salinity did not alter offspring reproduction timing in any F2 environment (Figure 3). Cumulative reproduction probabilities remained similar to those of control-maternal offspring, although reproduction tended to be lower early in life under the F2 elevated salinity environment, with credible intervals overlapping zero. Total offspring production followed a similar pattern, with offspring of salt-exposed mothers producing similar or marginally fewer offspring across environments (Figure 4). Thus, the strong survival benefits associated with maternal salinity did not translate into increased reproductive output.

Maternal salinity slightly reduced offspring body size under elevated salinity (day 16, –0.133, 90% CI –0.265 to –0.005) and combined warming–salinity conditions (–0.223, 90% CI –0.420 to –0.024; Figure 5). In contrast, body size remained unchanged under control and warming conditions.

## 4. Discussion

Our results show that environmental matching did not consistently predict whether maternal effects were beneficial. Maternal exposure to elevated salinity enhanced offspring survival under matching salinity, but these benefits extended beyond matching conditions to warming and multiple stressors. In contrast, maternal exposure to warming provided no detectable survival benefit under matching warming and imposed substantial survival costs in mismatching environments, particularly under elevated salinity. The consequences of maternal exposure also depended strongly on the fitness trait considered: maternal warming accelerated offspring reproduction across environments despite its costs to survival, whereas the survival benefits associated with maternal salinity were accompanied by reduced body size when offspring experienced elevated salinity and no increase in reproductive output. Together, these findings show that environmental matching alone provides an incomplete basis for predicting beneficial maternal effects, whose consequences depend on both the environmental context offspring encounter and the fitness trait through which they are expressed.

### 4.1 Environmental matching does not consistently predict beneficial maternal effects

Our results provide mixed support for the prediction that maternal exposure benefits offspring subsequently encountering the same environmental stressor. Maternal salinity substantially increased offspring survival under matching salinity, consistent with the prediction for an anticipatory maternal effect, whereas maternal warming provided no detectable survival advantage under matching warming. Such variation in the benefits of environmental matching has been observed across taxa. In *Daphnia*, parental warming can improve offspring performance under matching conditions, but evidence for adaptive thermal matching varies among studies, traits and environmental contexts (Betini *et al*. 2020; Kielland *et al*. 2017; Toyota *et al*. 2019; Walsh *et al*. 2014). Similarly, thermal matching in guppies (*Poecilia reticulata*) depends on the trait and number of generations exposed (Le Roy *et al*. 2017), while experimental evolution in the nematode *Caenorhabditis remanei* showed that anticipatory maternal effects evolved when parental and offspring environments were correlated but were lost when this predictability was removed (Lind *et al*. 2020). Differences in maternal condition are unlikely to fully explain our results: salinity-exposed F1 mothers did not consistently outperform control mothers (Fig. S1), despite their offspring showing enhanced survival across stressful environments, providing little support for a simple “silver-spoon” effect (Engqvist & Reinhold 2016).

Importantly, the benefits of maternal salinity extended beyond environmental matching, enhancing offspring survival under warming and, even more strongly, when warming and salinity occurred together. These broader benefits are consistent with generalized stress buffering or cross-tolerance, whereby maternal exposure to one stressor enhances offspring tolerance to another (Garbutt *et al*. 2014; Rodgers & Gomez Isaza 2021; Sun *et al*. 2023). Such buffering challenges a strict anticipatory-matching interpretation because maternal salinity was beneficial across several stressful offspring environments rather than specifically under the maternal environment. Conversely, maternal warming neither improved survival under matching warming nor provided broader survival benefits and instead imposed substantial costs under alternative environments. Environmental matching alone therefore cannot explain the magnitude or direction of maternal effects.

One explanation for these patterns is that maternal exposure does not necessarily induce offspring phenotypes that are specifically beneficial under the maternal environment. Different maternal stressors may instead affect distinct physiological, developmental or allocation pathways, generating offspring responses that differ in both their direction and environmental specificity (Donelson *et al*. 2018; Marshall 2008). Importantly, environmental matching and environmental predictability represent distinct concepts: matching describes whether maternal and offspring environments are the same, whereas predictability describes how reliably maternal conditions forecast the environments offspring are likely to encounter (Burgess & Marshall 2014; Kuijper & Hoyle 2015). Experimental evolution provides direct support for this distinction, with anticipatory maternal effects evolving when parental and offspring environments were correlated across generations but being lost when this predictability was removed (Lind *et al*. 2020). Our experiment manipulated matching rather than predictability and therefore cannot determine whether intergenerational predictability contributed to the contrasting effects of maternal salinity and warming. Explicitly manipulating predictability alongside environmental matching will be an important next step for determining when maternal effects represent environment-specific anticipatory responses and when they instead generate broader changes in offspring performance.

### 4.2 Multiple-stressor environments reveal context-dependent maternal effects

Multiple-stressor exposure revealed some of the most pronounced maternal effects in our experiment, supporting our prediction that their consequences would depend on the environmental context encountered by offspring. Maternal exposure to elevated salinity enhanced offspring survival when exposed to elevated salinity, warming and their combination, with a particularly pronounced difference under combined stressors. This larger difference should be interpreted cautiously because high survival under salinity alone may have limited the scope for additional maternal buffering. Nevertheless, the persistence of this benefit under combined warming and salinity shows that maternal effects remained important as offspring environmental complexity increased. Maternal exposure to warming produced a contrasting response: although it did not buffer survival under combined stressors, its positive effect on reproductive output was particularly pronounced in this environment. Thus, the same multiple-stressor environment revealed contrasting consequences of maternal history across traits, with maternal salinity associated primarily with enhanced survival and maternal warming with accelerated reproduction.

The pronounced maternal effects under combined warming and salinity may arise because maternal history alters how offspring cope with the overlapping physiological demands of thermal and osmotic stress. In *Daphnia*, salinity alters energy metabolism and osmoregulatory processes, including Na /K –ATPase activity, while combined salinity and warming can further modify metabolic and oxidative-stress responses (Garreta-Lara *et al*. 2018; Huang *et al*. 2026). Longer-term acclimation can also alter interactions between temperature and salinity, indicating that exposure history can modify sensitivity to their combined effects (Loureiro *et al*. 2015). Maternal exposure to elevated salinity may therefore have induced physiological adjustments that buffered some of the demands imposed by warming, but identifying the mechanisms will require direct measurements of osmoregulation, metabolism and oxidative stress.

These patterns highlight an important intersection between maternal effects and multiple-stressor ecology. Responses to multiple environmental drivers can differ substantially from those observed under either driver alone, limiting predictions based on isolated stressors (Cuenca Cambronero *et al*. 2021; Jackson *et al*. 2016; Orr *et al*. 2024). Empirical work in *Daphnia* further shows that transgenerational responses depend on stressor identity and combination, with maternal exposure to single versus combined stressors producing different consequences for offspring performance (Sun *et al*. 2023). Our results extend this context dependence across generations, showing that responses to interacting stressors depend not only on the conditions offspring experience, but also on maternal environmental history. Accounting for environmental history across generations may therefore be essential for predicting responses to interacting stressors, particularly when those responses differ among fitness traits.

### 4.3 Trait dependence changes what we classify as adaptive

Consistent with our third prediction, whether maternal effects appeared beneficial or detrimental depended strongly on the fitness trait considered. Maternal exposure to warming provides the clearest example: despite reducing offspring survival in several environments, it consistently accelerated reproduction, with particularly pronounced reproductive benefits under combined warming and salinity. This pattern is consistent with a shift toward a faster offspring life history rather than broadly increased stress tolerance. Similar responses have been observed in *Daphnia magna*, where mothers maintained at elevated temperature produced offspring that matured earlier and at smaller size and reproduced more frequently, without increasing lifetime reproductive success (Betini *et al*. 2020). Such shifts toward earlier reproduction may reflect greater investment in current reproduction when future reproductive opportunities are reduced, as predicted by life-history theory. Whether this represents an adaptive response, however, depends on whether early reproductive gains compensate for subsequent survival costs (Bell 1980; Duffield *et al*. 2017). The coexistence of accelerated reproduction and reduced survival following maternal warming therefore cautions against classifying maternal effects as adaptive from reproductive responses alone.

Maternal exposure to elevated salinity revealed a different trade-off. The survival benefits were not accompanied by increased reproduction and were associated with reduced body size when offspring experienced elevated salinity. Salinity can increase the energetic demands of osmoregulation in *Daphnia*, including Na /K –ATPase activity and metabolic rate, potentially reducing resources available for growth and reproduction (Smolders *et al*. 2005; Sun *et al*. 2024). Maternal salinity may therefore have shifted offspring allocation toward physiological maintenance and survival at the expense of growth, although the mechanisms remain to be tested. Together, these contrasting responses suggest that different maternal environments can generate distinct offspring life-history phenotypes: one characterized primarily by accelerated reproduction and the other by enhanced survival with potential growth costs.

Trait dependence therefore complicates the interpretation of adaptive transgenerational plasticity. Fitness emerges from the combined contributions of survival, growth and reproduction, yet maternal effects on these components need not align and may instead reflect underlying allocation trade-offs (Marks & Lailvaux 2024; Marshall 2008). Empirical studies similarly reveal divergent transgenerational responses among life-history traits across taxa (Betini *et al*. 2020; Coakley *et al*. 2018; Jarrold *et al*. 2019; Toyota *et al*. 2019), emphasizing that a benefit in one trait does not necessarily indicate an overall fitness advantage. Evaluating multiple fitness-related traits across offspring environments may therefore be essential for determining whether apparent benefits extend across fitness traits or instead reflect shifts in life-history allocation.

### 4.4 Conclusion: predicting maternal effects under environmental change

Predicting maternal effects under environmental change requires moving beyond maternal– offspring environmental matching to consider the ecological information provided by maternal environments, the combinations of stressors offspring encounter, and the fitness trait through which maternal effects are expressed. These dimensions may become increasingly important as environmental change alters environmental variability, temporal structure and stressor co-occurrence (Di Cecco & Gouhier 2018; Jackson *et al*. 2021). In particular, our findings highlight the need to extend multiple-stressor frameworks across generations, as responses to contemporary stressor combinations can depend on conditions experienced by previous generations (Tran *et al*. 2018). Multiple-stressor effects may therefore reflect not only the environmental drivers acting simultaneously on an organism, but also the sequence of conditions experienced across generations. Incorporating this temporal dimension will be important for predicting responses to environmental change, where the intensity, timing and co-occurrence of environmental stressors are themselves changing.

Our results show that maternal effects can extend beyond matching environments, persist under multiple stressors and generate contrasting consequences across fitness traits. A key next step will be to determine how these responses depend on the ecological information carried by maternal environments by experimentally manipulating intergenerational predictability alongside stressor identity and combination. Integrating such approaches with multiple fitness traits will help resolve when maternal effects buffer environmental change and when their benefits are offset by costs, improving our ability to predict their consequences in increasingly complex and variable environments.

## 6. Acknowledgements

The authors thank all laboratory members for their support with the empirical work. The authors thank Dieter Ebert, Michelle Krebs and Jürgen Hottinger for the provision of the *D. magna* genotype.

## Statement of authorship

C.C.B conceived the original research question, designed and conducted the experiment, analysed the data, and wrote the first draft of the manuscript. J.A.O developed and implemented the Bayesian statistical analyses and data analysis. F.V, P.L and J.P contributed to the development of the original research question. A.B, M.C, M.G, A.K and B.S contributed to conducting the experiment. All authors contributed to manuscript revision and approved the final version.

## Data accessibility statement

The data and code supporting the results of this study are publicly available in Zenodo at https://doi.org/10.5281/zenodo.23264479.

