## Supplementary material for "Beyond Environmental Matching: Maternal Effects Depend on Offspring Environmental Complexity and Fitness Traits"

**Running title:**  Maternal Effects Beyond Environmental Matching

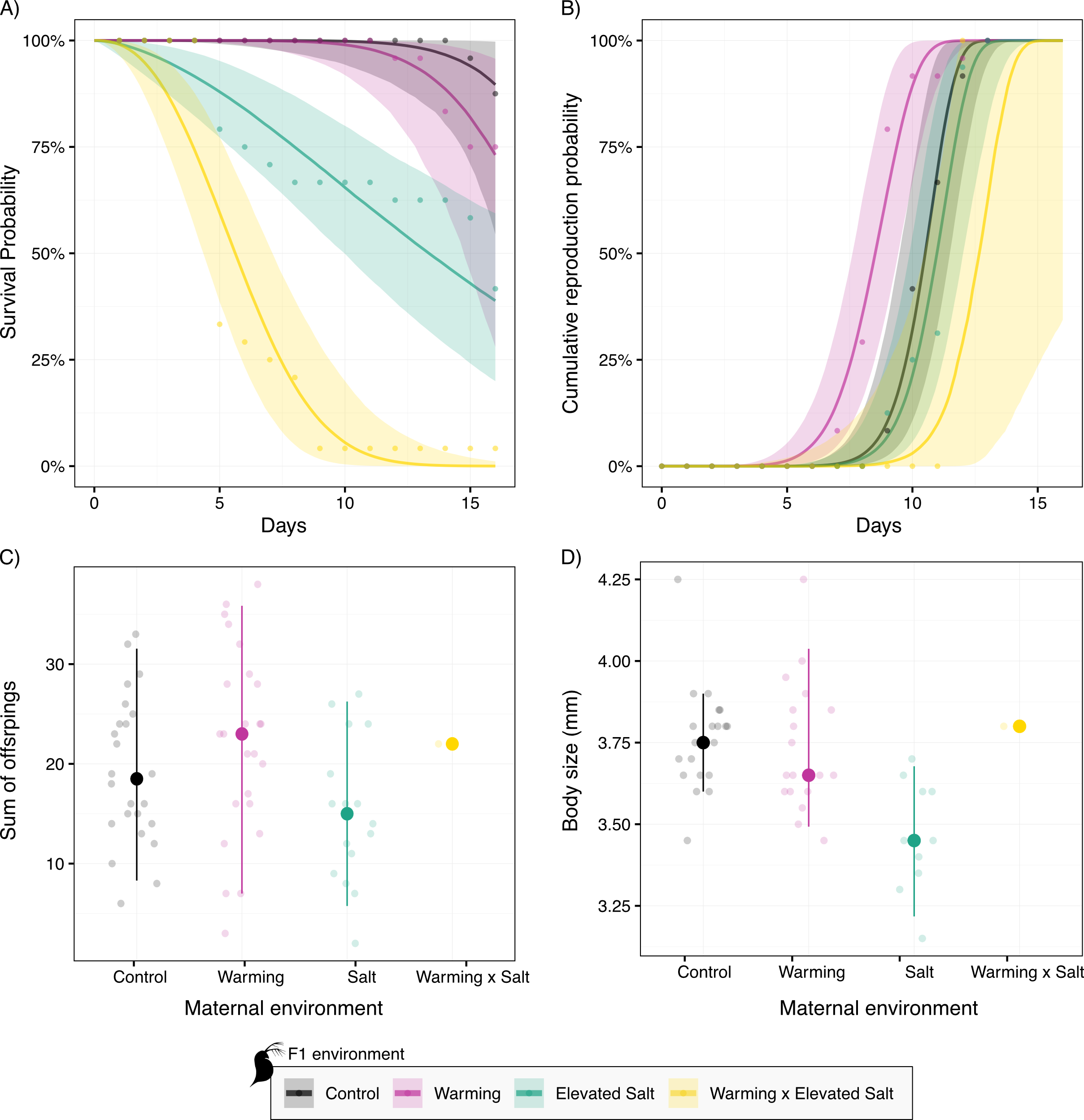

Figure S1: **Responses of F1 individuals to control conditions, warming, elevated salinity, and combined warming and elevated salinity.** Responses are shown for (A) survival, (B) cumulative reproduction probability, (C) total offspring number, and (D) body size of survivors. In panels A and B, lines represent median posterior predictions through time, shaded areas represent 90% credible intervals, and points show empirical observations. In panels C and D, filled circles represent median posterior predictions, lines represent 90% credible intervals, and shaded distributions represent the full posterior distributions.

Table S1: Realized temperature, salinity and conductivity across experimental treatments and generations. Values represent mean ± SD of measurements collected throughout each experimental phase. n indicates the number of measurements. Salinity of the control ADaM medium (0.33 g L⁻¹) represents the nominal concentration based on standardized medium preparation and was not measured directly.

| **Treatment** | **Generation** | **Target**  **temperature (°C)** | **Measured**  **temperature** | **Nominal**  **salinity (g L^-1^)** | **Measured**  **salinity (g L^-1^)** | **Conductivity**  (ms/cm) | **n** |
| --- | --- | --- | --- | --- | --- | --- | --- |
| Control | P | 20 | 20.1 ± 0.45 | 0.33 | - | 1.33 ± 0.04 | 36 |
| Control | F1 | 20 | 20.3 ± 0.67 | 0.33 | - | 1.33 ± 0.03 | 36 |
| Warming | F1 | 25 | 25.3 ± 0.72 | 0.33 | - | 1.33 ± 0.03 | 36 |
| Elevated salinity | F1 | 20 | 20.3 ± 0.67 | 5 | 5.07 ± 0.21 | 8.87 ± 0.11 | 36 |
| Combined | F1 | 25 | 25.3 ± 0.72 | 5 | 5.07 ± 0.21 | 8.87 ± 0.11 | 36 |
| Control | F2 | 20 | 20.2 ± 0.34 | 0.33 | - | 1.32 ± 0.05 | 36 |
| Warming | F2 | 25 | 24.9 ± 0.83 | 0.33 | - | 1.32 ± 0.05 | 36 |
| Elevated salinity | F2 | 20 | 20.2 ± 0.34 | 5 | 5.09 ± 0.25 | 8.92 ± 0.17 | 36 |
| Combined | F2 | 25 | 24.9 ± 0.83 | 5 | 5.09 ± 0.25 | 8.92 ± 0.17 | 36 |

Table S2: Posterior contrasts in offspring survival probability between offspring from mothers exposed to warming or elevated salinity and offspring from control mothers, calculated separately for each offspring environment and each day of the experiment. Values represent the posterior median difference in survival probability (treatment − control), with associated 90% credible intervals (CrIs). **% positive** and **% negative** indicate the percentage of posterior draws supporting positive or negative treatment effects, respectively. Positive median differences indicate higher survival of offspring from treated mothers relative to offspring from control mothers, whereas negative values indicate lower survival.

| **Maternal treatment** | **Offspring treatment** | **Day** | **Median difference** | **90% CrI** | **% positive** | **% negative** |
| --- | --- | --- | --- | --- | --- | --- |
| Warming | Control | 1 | -0.041 | -0.121 to 0.009 | 9.3 | 90.6 |
| Warming | Control | 2 | -0.075 | -0.176 to 0.003 | 5.5 | 94.5 |
| Warming | Control | 3 | -0.105 | -0.221 to -0.009 | 3.7 | 96.3 |
| Warming | Control | 4 | -0.132 | -0.262 to -0.021 | 2.5 | 97.5 |
| Warming | Control | 5 | -0.156 | -0.301 to -0.032 | 2.1 | 98.0 |
| Warming | Control | 6 | -0.179 | -0.341 to -0.042 | 1.7 | 98.4 |
| Warming | Control | 7 | -0.199 | -0.375 to -0.050 | 1.3 | 98.7 |
| Warming | Control | 8 | -0.217 | -0.407 to -0.057 | 1.1 | 98.9 |
| Warming | Control | 9 | -0.234 | -0.437 to -0.066 | 0.9 | 99.1 |
| Warming | Control | 10 | -0.248 | -0.466 to -0.072 | 0.8 | 99.2 |
| Warming | Control | 11 | -0.261 | -0.493 to -0.077 | 0.8 | 99.2 |
| Warming | Control | 12 | -0.274 | -0.514 to -0.081 | 0.8 | 99.2 |
| Warming | Control | 13 | -0.284 | -0.537 to -0.084 | 0.9 | 99.2 |
| Warming | Control | 14 | -0.296 | -0.554 to -0.088 | 0.8 | 99.2 |
| Warming | Control | 15 | -0.305 | -0.570 to -0.089 | 0.9 | 99.1 |
| Warming | Control | 16 | -0.313 | -0.582 to -0.090 | 0.9 | 99.1 |
| Warming | Warming | 1 | 0.033 | -0.067 to 0.131 | 71.7 | 28.3 |
| Warming | Warming | 2 | 0.046 | -0.075 to 0.165 | 74.1 | 25.9 |
| Warming | Warming | 3 | 0.055 | -0.081 to 0.186 | 75.0 | 25.0 |
| Warming | Warming | 4 | 0.061 | -0.087 to 0.205 | 75.3 | 24.7 |
| Warming | Warming | 5 | 0.066 | -0.091 to 0.225 | 75.4 | 24.6 |
| Warming | Warming | 6 | 0.069 | -0.097 to 0.239 | 74.9 | 25.1 |
| Warming | Warming | 7 | 0.071 | -0.104 to 0.252 | 74.7 | 25.4 |
| Warming | Warming | 8 | 0.074 | -0.111 to 0.263 | 73.8 | 26.2 |
| Warming | Warming | 9 | 0.076 | -0.120 to 0.276 | 73.5 | 26.5 |
| Warming | Warming | 10 | 0.079 | -0.126 to 0.286 | 73.3 | 26.7 |
| Warming | Warming | 11 | 0.080 | -0.131 to 0.294 | 72.8 | 27.2 |
| Warming | Warming | 12 | 0.080 | -0.140 to 0.304 | 72.1 | 27.9 |
| Warming | Warming | 13 | 0.080 | -0.144 to 0.311 | 71.7 | 28.3 |
| Warming | Warming | 14 | 0.080 | -0.151 to 0.316 | 71.0 | 28.9 |
| Warming | Warming | 15 | 0.080 | -0.158 to 0.323 | 70.5 | 29.5 |
| Warming | Warming | 16 | 0.079 | -0.166 to 0.329 | 70.0 | 30.0 |
| Warming | Salt | 1 | -0.057 | -0.150 to 0.003 | 5.9 | 94.2 |
| Warming | Salt | 2 | -0.106 | -0.226 to -0.018 | 2.4 | 97.7 |
| Warming | Salt | 3 | -0.151 | -0.291 to -0.041 | 1.1 | 98.9 |
| Warming | Salt | 4 | -0.192 | -0.349 to -0.066 | 0.5 | 99.5 |
| Warming | Salt | 5 | -0.229 | -0.400 to -0.089 | 0.4 | 99.6 |
| Warming | Salt | 6 | -0.263 | -0.446 to -0.110 | 0.3 | 99.7 |
| Warming | Salt | 7 | -0.292 | -0.486 to -0.128 | 0.2 | 99.8 |
| Warming | Salt | 8 | -0.319 | -0.523 to -0.144 | 0.1 | 99.9 |
| Warming | Salt | 9 | -0.342 | -0.554 to -0.158 | 0.1 | 99.9 |
| Warming | Salt | 10 | -0.366 | -0.585 to -0.169 | 0.1 | 99.9 |
| Warming | Salt | 11 | -0.386 | -0.611 to -0.178 | 0.1 | 99.9 |
| Warming | Salt | 12 | -0.405 | -0.631 to -0.188 | 0.1 | 99.9 |
| Warming | Salt | 13 | -0.422 | -0.651 to -0.197 | 0.1 | 99.9 |
| Warming | Salt | 14 | -0.436 | -0.666 to -0.203 | 0.1 | 99.9 |
| Warming | Salt | 15 | -0.449 | -0.681 to -0.210 | 0.1 | 99.9 |
| Warming | Salt | 16 | -0.462 | -0.692 to -0.214 | 0.1 | 99.9 |
| Warming | Warming_Salt | 1 | 0.021 | -0.104 to 0.141 | 60.9 | 39.1 |
| Warming | Warming_Salt | 2 | -0.000 | -0.159 to 0.149 | 50.0 | 50.0 |
| Warming | Warming_Salt | 3 | -0.029 | -0.207 to 0.144 | 39.6 | 60.4 |
| Warming | Warming_Salt | 4 | -0.055 | -0.248 to 0.136 | 32.1 | 67.9 |
| Warming | Warming_Salt | 5 | -0.076 | -0.278 to 0.127 | 26.9 | 73.2 |
| Warming | Warming_Salt | 6 | -0.095 | -0.308 to 0.120 | 23.5 | 76.5 |
| Warming | Warming_Salt | 7 | -0.110 | -0.332 to 0.115 | 21.6 | 78.4 |
| Warming | Warming_Salt | 8 | -0.123 | -0.347 to 0.110 | 19.9 | 80.1 |
| Warming | Warming_Salt | 9 | -0.132 | -0.361 to 0.109 | 18.7 | 81.3 |
| Warming | Warming_Salt | 10 | -0.139 | -0.369 to 0.108 | 17.8 | 82.2 |
| Warming | Warming_Salt | 11 | -0.144 | -0.372 to 0.102 | 17.0 | 83.0 |
| Warming | Warming_Salt | 12 | -0.145 | -0.378 to 0.097 | 16.3 | 83.7 |
| Warming | Warming_Salt | 13 | -0.144 | -0.377 to 0.093 | 16.2 | 83.8 |
| Warming | Warming_Salt | 14 | -0.142 | -0.377 to 0.089 | 16.2 | 83.8 |
| Warming | Warming_Salt | 15 | -0.142 | -0.377 to 0.087 | 15.9 | 84.1 |
| Warming | Warming_Salt | 16 | -0.138 | -0.375 to 0.083 | 15.8 | 84.2 |
| Salt | Control | 1 | 0.007 | -0.021 to 0.049 | 71.3 | 28.7 |
| Salt | Control | 2 | 0.014 | -0.027 to 0.074 | 74.7 | 25.4 |
| Salt | Control | 3 | 0.022 | -0.032 to 0.095 | 77.6 | 22.4 |
| Salt | Control | 4 | 0.030 | -0.033 to 0.112 | 79.8 | 20.2 |
| Salt | Control | 5 | 0.037 | -0.034 to 0.130 | 81.7 | 18.4 |
| Salt | Control | 6 | 0.045 | -0.034 to 0.144 | 83.0 | 17.0 |
| Salt | Control | 7 | 0.052 | -0.035 to 0.158 | 84.2 | 15.8 |
| Salt | Control | 8 | 0.059 | -0.036 to 0.172 | 85.1 | 14.9 |
| Salt | Control | 9 | 0.065 | -0.036 to 0.185 | 85.9 | 14.1 |
| Salt | Control | 10 | 0.072 | -0.035 to 0.198 | 86.5 | 13.5 |
| Salt | Control | 11 | 0.078 | -0.036 to 0.213 | 87.2 | 12.8 |
| Salt | Control | 12 | 0.085 | -0.036 to 0.227 | 87.7 | 12.3 |
| Salt | Control | 13 | 0.091 | -0.036 to 0.240 | 88.3 | 11.7 |
| Salt | Control | 14 | 0.098 | -0.036 to 0.255 | 88.6 | 11.4 |
| Salt | Control | 15 | 0.105 | -0.036 to 0.267 | 88.9 | 11.1 |
| Salt | Control | 16 | 0.110 | -0.035 to 0.281 | 89.3 | 10.7 |
| Salt | Warming | 1 | 0.102 | 0.029 to 0.194 | 98.9 | 1.1 |
| Salt | Warming | 2 | 0.152 | 0.058 to 0.261 | 99.6 | 0.4 |
| Salt | Warming | 3 | 0.189 | 0.084 to 0.313 | 99.8 | 0.2 |
| Salt | Warming | 4 | 0.220 | 0.104 to 0.358 | 99.9 | 0.1 |
| Salt | Warming | 5 | 0.246 | 0.119 to 0.397 | 99.9 | 0.1 |
| Salt | Warming | 6 | 0.267 | 0.131 to 0.432 | 99.9 | 0.1 |
| Salt | Warming | 7 | 0.285 | 0.143 to 0.461 | 99.9 | 0.1 |
| Salt | Warming | 8 | 0.302 | 0.150 to 0.486 | 99.9 | 0.1 |
| Salt | Warming | 9 | 0.314 | 0.156 to 0.510 | 99.9 | 0.1 |
| Salt | Warming | 10 | 0.327 | 0.160 to 0.531 | 99.9 | 0.1 |
| Salt | Warming | 11 | 0.337 | 0.164 to 0.549 | 99.9 | 0.1 |
| Salt | Warming | 12 | 0.346 | 0.166 to 0.567 | 99.9 | 0.1 |
| Salt | Warming | 13 | 0.355 | 0.167 to 0.584 | 99.9 | 0.1 |
| Salt | Warming | 14 | 0.363 | 0.166 to 0.598 | 99.8 | 0.2 |
| Salt | Warming | 15 | 0.371 | 0.167 to 0.612 | 99.8 | 0.2 |
| Salt | Warming | 16 | 0.378 | 0.167 to 0.621 | 99.7 | 0.3 |
| Salt | Salt | 1 | 0.019 | 0.002 to 0.065 | 98.8 | 1.2 |
| Salt | Salt | 2 | 0.034 | 0.006 to 0.097 | 99.1 | 0.9 |
| Salt | Salt | 3 | 0.047 | 0.010 to 0.123 | 99.2 | 0.8 |
| Salt | Salt | 4 | 0.060 | 0.015 to 0.145 | 99.2 | 0.8 |
| Salt | Salt | 5 | 0.073 | 0.020 to 0.166 | 99.4 | 0.7 |
| Salt | Salt | 6 | 0.084 | 0.025 to 0.186 | 99.4 | 0.7 |
| Salt | Salt | 7 | 0.095 | 0.029 to 0.205 | 99.4 | 0.6 |
| Salt | Salt | 8 | 0.107 | 0.034 to 0.220 | 99.5 | 0.5 |
| Salt | Salt | 9 | 0.118 | 0.038 to 0.239 | 99.5 | 0.5 |
| Salt | Salt | 10 | 0.128 | 0.042 to 0.255 | 99.5 | 0.5 |
| Salt | Salt | 11 | 0.138 | 0.047 to 0.273 | 99.5 | 0.5 |
| Salt | Salt | 12 | 0.147 | 0.051 to 0.289 | 99.6 | 0.4 |
| Salt | Salt | 13 | 0.156 | 0.054 to 0.303 | 99.6 | 0.4 |
| Salt | Salt | 14 | 0.165 | 0.058 to 0.319 | 99.6 | 0.4 |
| Salt | Salt | 15 | 0.174 | 0.062 to 0.336 | 99.6 | 0.4 |
| Salt | Salt | 16 | 0.182 | 0.065 to 0.348 | 99.5 | 0.5 |
| Salt | Warming_Salt | 1 | 0.140 | 0.056 to 0.246 | 99.9 | 0.1 |
| Salt | Warming_Salt | 2 | 0.220 | 0.108 to 0.354 | 100.0 | 0.0 |
| Salt | Warming_Salt | 3 | 0.280 | 0.148 to 0.438 | 100.0 | 0.0 |
| Salt | Warming_Salt | 4 | 0.327 | 0.178 to 0.502 | 100.0 | 0.0 |
| Salt | Warming_Salt | 5 | 0.365 | 0.202 to 0.555 | 100.0 | 0.0 |
| Salt | Warming_Salt | 6 | 0.396 | 0.221 to 0.597 | 100.0 | 0.0 |
| Salt | Warming_Salt | 7 | 0.421 | 0.236 to 0.629 | 100.0 | 0.0 |
| Salt | Warming_Salt | 8 | 0.440 | 0.242 to 0.653 | 100.0 | 0.1 |
| Salt | Warming_Salt | 9 | 0.455 | 0.244 to 0.669 | 99.9 | 0.1 |
| Salt | Warming_Salt | 10 | 0.465 | 0.244 to 0.683 | 99.9 | 0.1 |
| Salt | Warming_Salt | 11 | 0.473 | 0.243 to 0.694 | 99.9 | 0.1 |
| Salt | Warming_Salt | 12 | 0.478 | 0.235 to 0.703 | 99.9 | 0.1 |
| Salt | Warming_Salt | 13 | 0.481 | 0.223 to 0.710 | 99.9 | 0.1 |
| Salt | Warming_Salt | 14 | 0.480 | 0.210 to 0.714 | 99.7 | 0.4 |
| Salt | Warming_Salt | 15 | 0.479 | 0.194 to 0.717 | 99.5 | 0.5 |
| Salt | Warming_Salt | 16 | 0.475 | 0.175 to 0.716 | 99.3 | 0.7 |

Table S3: Posterior contrasts in offspring cumulative reproduction probability between offspring from mothers exposed to warming or elevated salinity and offspring from control mothers, calculated separately for each offspring environment and each day of the experiment. Values represent the posterior median difference in survival probability (treatment − control), with associated 90% credible intervals (CrIs). **% positive** and **% negative** indicate the percentage of posterior draws supporting positive or negative treatment effects, respectively. Positive median differences indicate earlier cumulative reproduction of offspring from treated mothers relative to offspring from control mothers, whereas negative values indicate a delayed reproduction. In

| **Maternal treatment** | **Offspring treatment** | **Day** | **Median difference** | **90% CrI** | **% positive** | **% negative** |
| --- | --- | --- | --- | --- | --- | --- |
| Warming | Control | 1 | 0.000 | -0.000 to 0.000 | 88.9 | 11.1 |
| Warming | Control | 2 | 0.000 | -0.000 to 0.001 | 90.2 | 9.8 |
| Warming | Control | 3 | 0.000 | -0.000 to 0.005 | 91.3 | 8.7 |
| Warming | Control | 4 | 0.001 | -0.000 to 0.019 | 93.0 | 7.0 |
| Warming | Control | 5 | 0.006 | -0.000 to 0.058 | 94.3 | 5.7 |
| Warming | Control | 6 | 0.023 | 0.000 to 0.136 | 95.5 | 4.5 |
| Warming | Control | 7 | 0.073 | 0.004 to 0.261 | 97.2 | 2.8 |
| Warming | Control | 8 | 0.185 | 0.027 to 0.443 | 98.5 | 1.5 |
| Warming | Control | 9 | 0.355 | 0.095 to 0.627 | 99.4 | 0.7 |
| Warming | Control | 10 | 0.478 | 0.178 to 0.722 | 99.6 | 0.4 |
| Warming | Control | 11 | 0.384 | 0.086 to 0.643 | 99.0 | 1.1 |
| Warming | Control | 12 | 0.126 | 0.001 to 0.430 | 96.6 | 3.3 |
| Warming | Control | 13 | 0.007 | -0.003 to 0.165 | 88.3 | 11.2 |
| Warming | Control | 14 | 0.000 | -0.001 to 0.026 | 74.8 | 22.4 |
| Warming | Control | 15 | 0.000 | -0.000 to 0.001 | 54.5 | 29.6 |
| Warming | Control | 16 | 0.000 | -0.000 to 0.000 | 30.0 | 27.1 |
| Warming | Warming | 1 | 0.000 | -0.000 to 0.001 | 92.2 | 7.8 |
| Warming | Warming | 2 | 0.001 | -0.000 to 0.012 | 93.3 | 6.7 |
| Warming | Warming | 3 | 0.006 | -0.000 to 0.049 | 94.3 | 5.7 |
| Warming | Warming | 4 | 0.027 | 0.000 to 0.129 | 95.5 | 4.5 |
| Warming | Warming | 5 | 0.084 | 0.005 to 0.261 | 96.5 | 3.5 |
| Warming | Warming | 6 | 0.200 | 0.033 to 0.443 | 97.7 | 2.4 |
| Warming | Warming | 7 | 0.364 | 0.098 to 0.630 | 98.8 | 1.2 |
| Warming | Warming | 8 | 0.482 | 0.137 to 0.753 | 98.8 | 1.2 |
| Warming | Warming | 9 | 0.404 | 0.037 to 0.729 | 97.1 | 2.9 |
| Warming | Warming | 10 | 0.159 | -0.022 to 0.558 | 90.1 | 9.9 |
| Warming | Warming | 11 | 0.015 | -0.027 to 0.309 | 76.5 | 23.4 |
| Warming | Warming | 12 | 0.000 | -0.013 to 0.123 | 61.8 | 37.3 |
| Warming | Warming | 13 | 0.000 | -0.004 to 0.031 | 48.8 | 45.0 |
| Warming | Warming | 14 | 0.000 | -0.001 to 0.005 | 36.9 | 44.0 |
| Warming | Warming | 15 | 0.000 | -0.000 to 0.000 | 26.9 | 38.1 |
| Warming | Warming | 16 | 0.000 | -0.000 to 0.000 | 18.6 | 29.8 |
| Warming | Salt | 1 | 0.000 | -0.000 to 0.003 | 94.2 | 5.8 |
| Warming | Salt | 2 | 0.002 | -0.000 to 0.018 | 94.2 | 5.8 |
| Warming | Salt | 3 | 0.007 | -0.000 to 0.049 | 94.1 | 5.9 |
| Warming | Salt | 4 | 0.020 | -0.000 to 0.100 | 94.1 | 5.9 |
| Warming | Salt | 5 | 0.044 | -0.002 to 0.166 | 93.8 | 6.2 |
| Warming | Salt | 6 | 0.079 | -0.004 to 0.245 | 93.4 | 6.6 |
| Warming | Salt | 7 | 0.123 | -0.012 to 0.330 | 92.7 | 7.3 |
| Warming | Salt | 8 | 0.165 | -0.029 to 0.407 | 91.5 | 8.5 |
| Warming | Salt | 9 | 0.192 | -0.057 to 0.455 | 89.8 | 10.2 |
| Warming | Salt | 10 | 0.187 | -0.107 to 0.460 | 85.5 | 14.4 |
| Warming | Salt | 11 | 0.139 | -0.162 to 0.413 | 78.3 | 21.7 |
| Warming | Salt | 12 | 0.067 | -0.214 to 0.313 | 66.4 | 33.6 |
| Warming | Salt | 13 | 0.007 | -0.235 to 0.206 | 52.8 | 47.2 |
| Warming | Salt | 14 | -0.014 | -0.222 to 0.113 | 39.8 | 60.2 |
| Warming | Salt | 15 | -0.014 | -0.188 to 0.049 | 28.8 | 71.2 |
| Warming | Salt | 16 | -0.007 | -0.143 to 0.016 | 22.7 | 77.3 |
| Warming | Warming_Salt | 1 | 0.000 | -0.000 to 0.009 | 76.0 | 23.9 |
| Warming | Warming_Salt | 2 | 0.001 | -0.001 to 0.041 | 79.5 | 20.5 |
| Warming | Warming_Salt | 3 | 0.007 | -0.005 to 0.095 | 82.8 | 17.2 |
| Warming | Warming_Salt | 4 | 0.028 | -0.010 to 0.180 | 87.1 | 13.0 |
| Warming | Warming_Salt | 5 | 0.091 | -0.009 to 0.317 | 92.3 | 7.6 |
| Warming | Warming_Salt | 6 | 0.218 | 0.002 to 0.537 | 95.2 | 4.8 |
| Warming | Warming_Salt | 7 | 0.410 | 0.061 to 0.827 | 97.5 | 2.5 |
| Warming | Warming_Salt | 8 | 0.592 | 0.085 to 0.936 | 97.4 | 2.6 |
| Warming | Warming_Salt | 9 | 0.597 | 0.054 to 0.917 | 96.5 | 3.5 |
| Warming | Warming_Salt | 10 | 0.459 | -0.040 to 0.822 | 93.8 | 6.2 |
| Warming | Warming_Salt | 11 | 0.252 | -0.137 to 0.656 | 89.5 | 10.4 |
| Warming | Warming_Salt | 12 | 0.082 | -0.198 to 0.443 | 84.3 | 15.7 |
| Warming | Warming_Salt | 13 | 0.011 | -0.214 to 0.274 | 78.7 | 21.1 |
| Warming | Warming_Salt | 14 | 0.000 | -0.191 to 0.150 | 73.1 | 25.1 |
| Warming | Warming_Salt | 15 | 0.000 | -0.157 to 0.077 | 65.1 | 27.9 |
| Warming | Warming_Salt | 16 | 0.000 | -0.127 to 0.032 | 55.6 | 29.3 |
| Salt | Control | 1 | -0.000 | -0.000 to 0.000 | 49.2 | 50.3 |
| Salt | Control | 2 | -0.000 | -0.000 to 0.000 | 49.9 | 50.1 |
| Salt | Control | 3 | 0.000 | -0.000 to 0.000 | 50.3 | 49.7 |
| Salt | Control | 4 | 0.000 | -0.000 to 0.000 | 50.9 | 49.1 |
| Salt | Control | 5 | 0.000 | -0.002 to 0.002 | 51.4 | 48.6 |
| Salt | Control | 6 | 0.000 | -0.009 to 0.009 | 51.9 | 48.1 |
| Salt | Control | 7 | 0.000 | -0.027 to 0.029 | 52.5 | 47.4 |
| Salt | Control | 8 | 0.002 | -0.067 to 0.078 | 54.0 | 46.0 |
| Salt | Control | 9 | 0.011 | -0.133 to 0.165 | 56.5 | 43.5 |
| Salt | Control | 10 | 0.032 | -0.191 to 0.272 | 60.2 | 39.9 |
| Salt | Control | 11 | 0.055 | -0.187 to 0.312 | 64.7 | 35.3 |
| Salt | Control | 12 | 0.032 | -0.109 to 0.233 | 68.9 | 30.9 |
| Salt | Control | 13 | 0.002 | -0.037 to 0.097 | 67.5 | 32.1 |
| Salt | Control | 14 | 0.000 | -0.004 to 0.018 | 62.7 | 35.1 |
| Salt | Control | 15 | 0.000 | -0.000 to 0.001 | 52.4 | 32.2 |
| Salt | Control | 16 | 0.000 | -0.000 to 0.000 | 31.9 | 20.6 |
| Salt | Warming | 1 | -0.000 | -0.000 to 0.000 | 40.7 | 59.2 |
| Salt | Warming | 2 | -0.000 | -0.000 to 0.000 | 42.4 | 57.6 |
| Salt | Warming | 3 | -0.000 | -0.004 to 0.001 | 44.7 | 55.3 |
| Salt | Warming | 4 | -0.000 | -0.015 to 0.005 | 47.0 | 53.0 |
| Salt | Warming | 5 | 0.000 | -0.044 to 0.023 | 50.4 | 49.6 |
| Salt | Warming | 6 | 0.003 | -0.098 to 0.077 | 55.1 | 44.9 |
| Salt | Warming | 7 | 0.026 | -0.166 to 0.206 | 63.0 | 37.0 |
| Salt | Warming | 8 | 0.120 | -0.190 to 0.415 | 75.4 | 24.6 |
| Salt | Warming | 9 | 0.231 | -0.119 to 0.575 | 86.9 | 13.1 |
| Salt | Warming | 10 | 0.143 | -0.030 to 0.518 | 89.8 | 10.2 |
| Salt | Warming | 11 | 0.018 | -0.002 to 0.320 | 87.7 | 11.9 |
| Salt | Warming | 12 | 0.000 | -0.000 to 0.126 | 82.2 | 14.4 |
| Salt | Warming | 13 | 0.000 | -0.000 to 0.033 | 70.7 | 12.2 |
| Salt | Warming | 14 | 0.000 | -0.000 to 0.005 | 51.8 | 6.6 |
| Salt | Warming | 15 | 0.000 | 0.000 to 0.000 | 34.9 | 3.0 |
| Salt | Warming | 16 | 0.000 | 0.000 to 0.000 | 22.6 | 1.5 |
| Salt | Salt | 1 | -0.000 | -0.000 to 0.000 | 18.1 | 82.0 |
| Salt | Salt | 2 | -0.000 | -0.000 to 0.000 | 18.1 | 81.9 |
| Salt | Salt | 3 | -0.000 | -0.003 to 0.000 | 18.1 | 81.9 |
| Salt | Salt | 4 | -0.001 | -0.010 to 0.002 | 17.8 | 82.2 |
| Salt | Salt | 5 | -0.004 | -0.027 to 0.005 | 18.0 | 82.0 |
| Salt | Salt | 6 | -0.011 | -0.059 to 0.013 | 18.2 | 81.8 |
| Salt | Salt | 7 | -0.025 | -0.112 to 0.028 | 18.4 | 81.7 |
| Salt | Salt | 8 | -0.049 | -0.182 to 0.050 | 18.9 | 81.1 |
| Salt | Salt | 9 | -0.081 | -0.258 to 0.079 | 19.6 | 80.4 |
| Salt | Salt | 10 | -0.103 | -0.323 to 0.114 | 21.3 | 78.7 |
| Salt | Salt | 11 | -0.098 | -0.339 to 0.145 | 25.1 | 74.9 |
| Salt | Salt | 12 | -0.056 | -0.294 to 0.163 | 33.3 | 66.7 |
| Salt | Salt | 13 | -0.008 | -0.202 to 0.145 | 46.4 | 53.6 |
| Salt | Salt | 14 | 0.005 | -0.100 to 0.112 | 58.4 | 41.6 |
| Salt | Salt | 15 | 0.003 | -0.036 to 0.065 | 67.8 | 32.2 |
| Salt | Salt | 16 | 0.000 | -0.008 to 0.029 | 72.2 | 27.6 |
| Salt | Warming_Salt | 1 | 0.000 | -0.000 to 0.000 | 70.8 | 29.2 |
| Salt | Warming_Salt | 2 | 0.000 | -0.003 to 0.003 | 71.1 | 28.9 |
| Salt | Warming_Salt | 3 | 0.001 | -0.013 to 0.012 | 71.4 | 28.6 |
| Salt | Warming_Salt | 4 | 0.003 | -0.033 to 0.033 | 72.0 | 28.0 |
| Salt | Warming_Salt | 5 | 0.010 | -0.065 to 0.072 | 72.7 | 27.3 |
| Salt | Warming_Salt | 6 | 0.026 | -0.107 to 0.134 | 73.4 | 26.6 |
| Salt | Warming_Salt | 7 | 0.055 | -0.154 to 0.222 | 74.2 | 25.8 |
| Salt | Warming_Salt | 8 | 0.096 | -0.193 to 0.324 | 75.0 | 25.0 |
| Salt | Warming_Salt | 9 | 0.130 | -0.218 to 0.412 | 75.0 | 24.9 |
| Salt | Warming_Salt | 10 | 0.129 | -0.228 to 0.451 | 72.2 | 27.8 |
| Salt | Warming_Salt | 11 | 0.075 | -0.216 to 0.414 | 66.0 | 34.1 |
| Salt | Warming_Salt | 12 | 0.016 | -0.175 to 0.343 | 56.3 | 43.7 |
| Salt | Warming_Salt | 13 | -0.000 | -0.117 to 0.237 | 48.1 | 51.8 |
| Salt | Warming_Salt | 14 | -0.000 | -0.059 to 0.144 | 43.0 | 56.8 |
| Salt | Warming_Salt | 15 | -0.000 | -0.024 to 0.081 | 39.9 | 59.4 |
| Salt | Warming_Salt | 16 | -0.000 | -0.008 to 0.039 | 38.3 | 59.5 |

Table S4. Posterior contrasts in total offspring production at the end of the experiment (day 16) between offspring from mothers exposed to warming or elevated salinity and offspring from control mothers, calculated separately for each offspring environment. Values represent the posterior median difference in total offspring production (treatment − control), with associated 90% credible intervals (CrIs). **% positive** and **% negative** indicate the percentage of posterior draws supporting positive or negative treatment effects, respectively. Positive median differences indicate greater total offspring production by offspring from treated mothers relative to offspring from control mothers, whereas negative values indicate lower total offspring production.

| **Maternal treatment** | **Offspring treatment** | **Day** | **Median difference** | **90% CrI** | **% positive** | **% negative** |
| --- | --- | --- | --- | --- | --- | --- |
| Warming | Control | 16 | 2.910 | -0.161 to 6.280 | 94.1 | 5.9 |
| Warming | Warming | 16 | 5.164 | -0.100 to 10.472 | 94.6 | 5.4 |
| Warming | Salt | 16 | -0.886 | -3.146 to 1.547 | 25.9 | 74.2 |
| Warming | Warming_Salt | 16 | 15.683 | 9.326 to 23.313 | 100.0 | 0.0 |
| Salt | Control | 16 | 1.426 | -1.125 to 4.201 | 82.5 | 17.5 |
| Salt | Warming | 16 | 1.884 | -2.968 to 6.433 | 73.7 | 26.4 |
| Salt | Salt | 16 | -1.468 | -3.365 to 0.302 | 8.6 | 91.5 |
| Salt | Warming_Salt | 16 | -2.005 | -5.643 to 1.280 | 17.0 | 83.0 |

Table S5. Posterior contrasts in offspring body size at the end of the experiment (day 16) between offspring from mothers exposed to warming or elevated salinity and offspring from control mothers, calculated separately for each offspring environment. Values represent the posterior median difference in body size (treatment − control), with associated 90% credible intervals (CrIs). **% positive** and **% negative** indicate the percentage of posterior draws supporting positive or negative treatment effects, respectively. Positive median differences indicate larger offspring body size in offspring from treated mothers relative to offspring from control mothers, whereas negative values indicate smaller offspring body size.

| **Maternal treatment** | **Offspring treatment** | **Day** | **Median difference** | **90% CrI** | **% positive** | **% negative** |
| --- | --- | --- | --- | --- | --- | --- |
| Warming | Control | 16 | -0.129 | -0.295 to 0.041 | 10.6 | 89.4 |
| Warming | Warming | 16 | 0.078 | -0.147 to 0.291 | 71.3 | 28.7 |
| Warming | Salt | 16 | 0.027 | -0.155 to 0.206 | 59.1 | 40.9 |
| Warming | Warming_Salt | 16 | 0.173 | -0.103 to 0.441 | 85.4 | 14.6 |
| Salt | Control | 16 | -0.119 | -0.261 to 0.012 | 6.6 | 93.4 |
| Salt | Warming | 16 | -0.044 | -0.247 to 0.159 | 35.8 | 64.2 |
| Salt | Salt | 16 | -0.133 | -0.265 to -0.005 | 4.4 | 95.6 |
| Salt | Warming_Salt | 16 | -0.223 | -0.420 to -0.024 | 3.5 | 96.5 |
